# Chromosome-scale genome assembly for Yellow Wood sorrel, *Oxalis stricta*

**DOI:** 10.1101/2025.06.26.661616

**Authors:** Joshua C. Wood, John P. Hamilton, Brieanne Vaillancourt, Julia Brose, Patrick P. Edger, C. Robin Buell

## Abstract

Yellow wood sorrel (*Oxalis stricta* L.), also known as sourgrass, juicy fruit, or sheep weed, is a member of the Oxalidaceae family. Yellow wood sorrel is commonly considered a weed and while native to North America, it is distributed across Europe, Asia, and Africa. To date, only two other genomes from the Oxalidaceae family have been published, star fruit (*Averrhoa carambola* L.) and *Oxalis articulata* Savingy. Here, we present a chromosome-scale assembly for *O. stricta*, revealing its allotetraploid nature and synteny within its two subgenomes as well as synteny with *A. carambola* and *O. articulata*. Using Oxford Nanopore Technologies long-read sequences coupled with chromatin capture sequencing, we generated a 436 Mb chromosome-scale assembly of *O. stricta* with a scaffold N50 length of 36.2 Mb that is anchored to 12 chromosomes across the two subgenomes. Assessment of the final genome assembly using the Long Terminal Repeat Assembly Index yielded a score of 13.12 and assessment of Benchmarking Universal Single Copy Orthologs revealed 99.3% complete orthologs; both metrics are suggestive of a high-quality reference genome. Total repetitive sequence content in the *O. stricta* genome was 39.7% with retroelements being the largest class of transposable elements. Annotation of protein-coding genes yielded 61,550 high confidence genes encoding 115,089 gene models. Synteny between the two *O. stricta* subgenomes was present in 91 syntenic blocks containing 40,705 genes, of which, 76.6% were present in 1:1 syntenic relationships between the two subgenomes. The availability of an annotated chromosome-scale high quality genome assembly for *O. stricta* will provide a launching point to understand the high fecundity of this weed and provide further foundation for comparative genomics within the Oxalidaceae.

## INTRODUCTION

*Oxalis stricta* L., also known as yellow wood sorrel, is a pervasive species in North America and is considered a weed (Holt and Elmore 1985)(Fig. 1). Yellow wood sorrel belongs to the Oxalidaceae, a name that aptly reflects the plants’ production of oxalic acid. The Oxalidaceae contains numerous species of botanical interest, including star fruit (*Averrhoa carambola* L.) and oca (*Oxalis tuberosa* Molina), a tuberous food crop in South America, as well as the ornamentals *Oxalis triangularis* A. St.-Hil. (False Shamrock) and *Oxalis versicolor* L. (Candy Cane Sorrel). To date, the only species from Oxalidaceae with a genome sequence are star fruit and *Oxalis articulata* Savingy (Wu *et al*. 2020; Yang *et al*. 2025).

**Figure 1:**
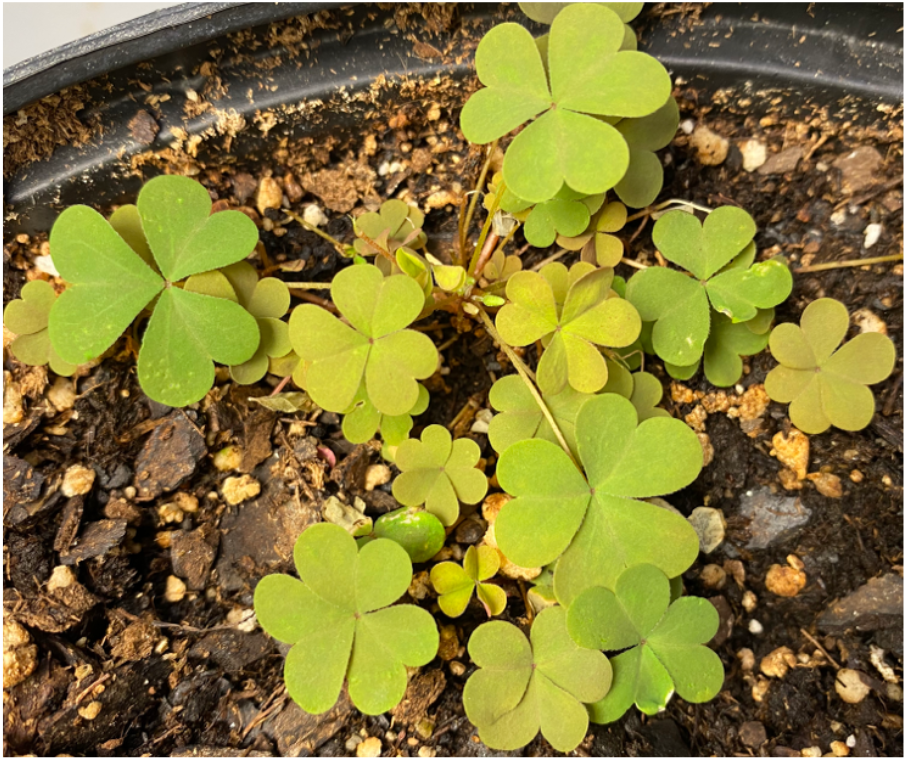
Picture of woodsorrel, *Oxalis stricta*

Yellow wood sorrel boasts edible leaves and seed pods containing oxalic acid and high levels of vitamin C, which impart a strong, tangy flavoring that contributes to its use as an herb (Shad *et al*. 2013). Despite these potential culinary applications, yellow wood sorrel is widely considered an invasive weed due to several key characteristics: its high fecundity, with each plant producing numerous seeds (Stevens 1932); the ability of its seed capsules to explosively disperse seeds up to 2 meters (van der Pijl 2012); the absence of a significant period of seed dormancy allowing for rapid germination; and its production of rhizomes, enabling perennial growth and persistence (Marshall 1987; Christopher Marble 2018).

*O. stricta* is a tetraploid with a base chromosome number of 6 (2n = 4x = 24) with reported 1C genome sizes that range between 528 Mb (Vaio *et al*. 2013) and 547 Mb (Bai *et al*. 2012). In this study, we generated a chromosome-scale genome assembly for *O. stricta* comprised of twelve chromosomes, spanning both two subgenomes, with a total assembly size of 436 Mb. We annotated 61,550 representative high-confidence genes encoding 115,089 gene models. This genomic resource will facilitate research within the Oxalidaceae, which contain oxalic acid content as well as the presence and absence of below ground storage organ formation.

## MATERIALS & METHODS

### Genomic DNA isolation and sequencing

Seeds of *O. stricta* were obtained from EdenWilds (Brooklyn, NY USA) and subjected to single seed descent. Plants were grown in a growth chamber under 21.1°C day/15.6°C night with 500 μmol/m^2^/s light intensity under a 15-hour photoperiod. DNA for short read sequencing was isolated from mature leaves using the Qiagen DNeasy Plant Pro Kit (Germantown, MD) and a library constructed using the PerkinElmer NEXTFLEX Rapid XP DNA-Seq Kit HT with NEXTFLEX UDI Barcodes (PerkinElmer, Waltham, MA). Sequencing was performed on an Illumina NovaSeq 6000 (San Diego, CA) generating paired end 150 nt reads (Illumina, San Diego, CA; Supplementary Table 1). Library preparation and sequencing was performed by the Texas A&M AgriLife Research: Genomics and Bioinformatics Service.

Plants were dark-treated for 24 hours prior to harvesting of leaf tissue for high molecular weight (HMW) DNA isolation as described previously (Vaillancourt and Buell 2019). HMW DNA was input into the Oxford Nanopore Technologies (ONT) Ligation sequencing gDNA kit (SQK-LSK110) and the resulting libraries sequenced on R9 FLO-MIN106 Rev D flow cells on a MinION (ONT, Oxford, UK; Supplementary Table 1). Young leaves were collected from plants grown in a greenhouse at 25°C day/15°C night, with 300 μmol/m^2^/s light intensity under 15 hours of light. Tissue was then used to construct two Hi-C libraries using the Proximo Plant v4.0 protocol with a modified fragmentation enzyme cocktail containing *Dpn*II, *Dde*I, *Hinf*I, and *Mse*I (Phase Genomics, Seattle, WA). Sequencing was performed on an Illumina NovaSeq 6000 at the Michigan State University Research Technology Support Facility generating paired end 150 nt reads (Supplementary Table 1).

### RNA isolation and sequencing

For gene expression abundance estimation and genome annotation, we constructed a replicated developmental tissue atlas that included a set of abiotic and biotic stresses to fully capture the *O. stricta* transcriptome. Plants were grown in a growth chamber under 21.1°C day/15.6°C night with 500 μmol/m^2^/s light intensity under a 15-hour photoperiod. Core developmental tissues included stem, flower (closed bud, open bud), fibrous root, as well as leaf tissue from a 24-hour diurnal time-course sampled every four hours (Supplementary Table 1). To mimic stress conditions, leaves were treated with Methyl Jasmonate (MeJA, 250μM) and benzothiadiazole (BTH, 100μg/ml) via leaf drenches with collection occurring 24 hrs after treatment. For salt treatment, 100mL of 150mM NaCl solution was applied to the pots and samples collected after 24 hrs. For cold treatment, plants were subjugated to a constant temperature of 10°C for 24 hours prior to collection. For heat treatment, plants were watered well to avoid drought stress and the temperature increased to 37°C (day) and 28°C (night) and samples collected after 24 hours. Finally, a drought treatment was conducted with leaves being collected after visible wilting was observed.

RNA was isolated using a modified hot borate method (Wan and Wilkins 1994) and residual DNA removed using the TURBO DNase kit (Invitrogen, Waltham, MA). RNA-Seq libraries were constructed using NEXTFLEX Poly(A) Beads 2.0 for PolyA selection followed by the PerkinElmer NEXTFLEX Rapid Directional RNA-Seq Kit 2.0 with NEXTFLEX RNA-Seq 2.0 Unique Dual Index Barcodes (PerkinElmer, Waltham, MA). Sequencing was performed on an Illumina NovaSeq 6000 (Illumina, San Diego, CA) generating paired end 150 nt reads. Library preparation and sequencing was performed by the Texas A&M AgriLife Research: Genomics and Bioinformatics Service. Full-length cDNA libraries were constructed using the ONT PCR-cDNA Barcoding kit (SQK-PCB109) and sequenced on a MinION using R9 FLO-MIN106 Rev D flow cells (ONT, Oxford, UK; Supplementary Table 1).

### Genome Assembly

The ONT genomic data was base-called using Guppy (v5.0.14) (https://nanoporetech.com/software/other/guppy/history?version=5-0-14) using the high accuracy model (dna_r9.4.1_450bps_hac.cfg) and the parameters --trim_strategy dna and --calib_detect. Reads shorter than 10kb were removed from the read pool using seqkit (v0.16.1) (Shen *et al*. 2024). Reads 10kb or greater were input into the Flye (v2.9) assembler (Kolmogorov *et al*. 2019) with options --nano-raw, --iterations 0, and --scaffold. The assembly was then polished using two rounds of Racon (v1.4.20) (Vaser *et al*. 2017) with the parameters set as follows: -m 8 -x -6 -g -8 -w 500 -u. After Racon, two rounds of Medaka (v1.4.4) (https://github.com/nanoporetech/medaka) polishing was completed using the basecaller model r941_min_hac_g507. Pilon (v1.24) (Walker *et al*. 2014) was then run using the Illumina whole genome shotgun reads in three consecutive rounds using the options --frags and --fix bases. Pseudomolecules were constructed using the Hi-C libraries as input into Juicer (v1.6) (Durand *et al*. 2016b) and 3D-DNA (v180922) (Dudchenko *et al*. 2017) with options: -i 10000 and -r 5. Juicebox (Durand *et al*. 2016b) was used to complete manual curation and the creation of the final chromosomes. The assembly was then filtered to remove contigs less than 10kb using seqkit (v0.16.1).

Multiple methods were used to assess the quality and completeness of the final assembly. First, the KAT software (v2.4.2) (Mapleson *et al*. 2017) was used to assess completeness of the final assembly based on k-mer representation in the final assembly. Second, the completeness of the final genome was assessed by determining the fraction of Benchmarking Single Copy Orthologs (BUSCO) using BUSCO v5 (Manni *et al*. 2021) (*Embryophyta* odb 10). Third, the Long Terminal Repeat (LTR) Assembly Index (LAI) metric (Ou *et al*. 2018) was used to determine contiguity of LTRs. Intact LTR-RTs (LTR retrotransposons) were identified using LTRharvest (v1.6.2) (Ellinghaus *et al*. 2008), LTR_FINDER_parallel (v1.0.7) (Ou and Jiang 2019), and LTR_retriever (v2.9.0) (Ou and Jiang 2018). Options for LTRharvest were: -minlenltr 100 - maxlenltr 7000 -mintsd 4 -maxtsd 6 -motif TGCA -motifmis 1 -similar 85 -vic 10 -seed 20. The options for LTR_FINDER_parallel were: -size 1000000 -time 300.

### Genome Annotation

The genome assembly was repeat masked by first creating a custom repeat library (CRL) for the genome. Repeats were first identified using RepeatModeler (v2.03) (Flynn *et al*. 2020) with protein-coding genes filtered out from the repeat database using ProtExcluder (v1.2) (Campbell *et al*. 2014) to create a CRL. The CRL was then combined with Viridiplantae repeats from RepBase (v20150807) (Bao *et al*. 2015) to generate the final CRL. The genome assembly was repeat-masked using the final CRL using RepeatMasker (v4.1.2-p1) (Chen 2004) with the parameters -e ncbi -s -nolow -no_is -gff.

RNA-seq libraries were processed for genome annotation by first cleaning with Cutadapt (v2.10) (Martin 2011) using a minimum length of 100 nt and quality cutoff of 10 then aligning the cleaned reads to the genome assembly using HISAT2 (2.1.0) (Kim *et al*. 2019). Oxford Nanopore (ONT) cDNA reads were processed with Pychopper (v2.5.0) (https://github.com/epi2me-labs/pychopper) and trimmed reads greater than 500 nt were aligned to the genome assembly using minimap2 (v2.17-r941) (Li 2021) with a maximum intron length of 5,000 nt. The aligned RNA-seq and ONT cDNA reads were each assembled using Stringtie (v2.2.1) (Kovaka *et al*. 2019) and transcripts less then 500 nt were removed.

Initial gene models were created using BRAKER2 (v2.1.6) (Brůna *et al*. 2021) using the soft-masked genome assembly and the aligned RNA-seq libraries as hints. The gene models were then refined using two rounds of PASA2 (v2.5.2) (Haas *et al*. 2003) to create a working gene model set. High-confidence gene models were identified from the working gene model set by filtering out gene models without expression evidence, or a PFAM domain match, or were a partial gene mode, or contained an interior stop codon. Functional annotation was assigned by searching the working gene models proteins against the TAIR (v10) (Lamesch *et al*. 2012) database and the Swiss-Prot plant proteins (release 2015_08) database using BLASTP (v2.12.0) (Altschul *et al*. 1990) and the PFAM (v35.0) (Mistry *et al*. 2021) database using PfamScan (v1.6) and assigning the annotation based on the first significant hit. Transposable elements were annotated using Extensive *de-novo* TE Annotator (EDTA) (v2.2.0) (Ou *et al*. 2019) with the genome and the working gene model CDS sequences as input.

### Assigning subgenomes

To define the subgenomes, annotated genes were aligned with GMAP (v2021-08-25) (Wu and Watanabe 2005) to the *A. carambola* genome (Wu *et al*. 2020). For each pair of chromosomes, the chromosome with the higher alignment rate to star fruit was binned into the “A” subgenome and the other chromosome into the “B” subgenome.

### Gene expression analyses

RNA-seq reads were checked for quality with FastQC (v0.11.9) (https://www.bioinformatics. babra-ham. ac.uk/projects/fastqc *FastQC*) and MultiQC (v1.11) (Ewels *et al*. 2016) before cleaning with Cutadapt (v4.4) using the options -q 30, -m 40, --trim-n, and -n 2. Reads were checked for contaminants using Kraken2 (v2.1) (Wood *et al*. 2019) with the database k2_pluspfp_20220908. Reads were then subsampled using seqtk (v1.3) (https://github.com/lh3/seqtk.git) to the median number of total reads after cleaning (65 M reads). To determine expression abundances, the quant algorithm of kallisto (v0.48.0) (Bray *et al*. 2016) was used along with the high confidence representative gene models with the stranded option set (−-rf-stranded) and the k-mer size at the default of 31.

### Comparative genome analyses

GENESPACE (v1.2.3) (Lovell *et al*. 2022) was used to identify syntelogs between *A. carambola* (Wu *et al*. 2020), *O. articulata* (Lovell *et al*. 2022), and *O. stricta*. The syntenic orthologs, hereafter ‘syntelogs’, were extracted using the ‘PASS’ flag from the *query*_pangenes function. OrthoFinder2 (v2.5.4) (Emms and Kelly 2019) was used to generate orthologous and paralogous groups using the predicted proteomes of *Arabidopsis thaliana* (Cheng *et al*. 2017), *A. carambola* (Wu *et al*. 2020), *Cephalotus follicularis* (Fukushima *et al*. 2017), *Oryza sativa* (Kawahara *et al*. 2013), *O. articulata* (Yang *et al*. 2025), *O. stricta*, and *Solanum tuberosum* (Pham *et al*. 2020). Gene Ontology (GO) associations were generated using InterProScan (v 5.69-101.0) (Jones *et al*. 2014) and enrichment of GO associations determined using TopGO (2.56.0) (Alexa *et al*. 2006).

## RESULTS & DISCUSSION

### Assembly of the *O. stricta* genome

A total of 44.9 Gb of genomic ONT long reads with an N50 read length of 37.2 kb (∼ 83x coverage) were used to assemble the *O. stricta* genome using the Flye genome assembler software (Supplementary Table 2). Following polishing with ONT long reads and Illumina short reads and scaffolding using Hi-C reads (Fig. 2), the *O. stricta* genome assembly totaled 436,804,292 bp present on 99 scaffolds with an N50 scaffold length of 36,226,067 bp (Table 1; Supplementary Table 3). Only 36 gaps are present within the assembly, totaling 18,036 bp of sequence. The 12 chromosomes represent 433,794,823 bp of sequence with the remaining 3,009,469 bp of unanchored sequence present on the remaining 87 scaffolds, resulting in a high level of continuity (Supplementary Table 4). KAT also revealed substantial representation of k-mers as single copy in the final assembly (Supplementary Figure 1). The LAI assays for intact LTR retrotransposons (LTR-RTs) to evaluate assembly continuity with a higher LAI score indicative of a more complete genome due to a greater number of intact LTR-RTs. The *O. stricta* LAI score was 13.12, placing the assembly within the category of a ‘reference quality’ genome (Ou *et al*. 2018). Assessment of genome completeness using BUSCO revealed 99.4% complete BUSCOs with a majority of orthologs duplicated suggesting the presence of subgenomes in the assembly (Supplementary Table 5), consistent with the reported tetraploidy of *O. stricta* (Vaio *et al*. 2013). Overall, these metrics indicate a high quality, chromosome-scale reference genome assembly for *O. stricta*.

**Table 1:** Assembly statistics for the *Oxalis stricta* assembly

| Species | Number of Sequences | Total Assembly Size (bp) | Min Scaffold Length (bp) | Max Scaffold Length (bp) | N50 Scaffold Length (bp) |
| --- | --- | --- | --- | --- | --- |
| *O. stricta* (Subgenome A) | 6 | 217,252,839 | 26,919,296 | 46,093,068 | 36,226,067 |
| *O. stricta* (Subgenome B) | 6 | 216,541,984 | 27,369,382 | 46,270,890 | 36,921,660 |
| *O. stricta* (v1) | 99 | 436,804,292 | 10,023 | 46,270,890 | 36,226,067 |

**Figure 2:**
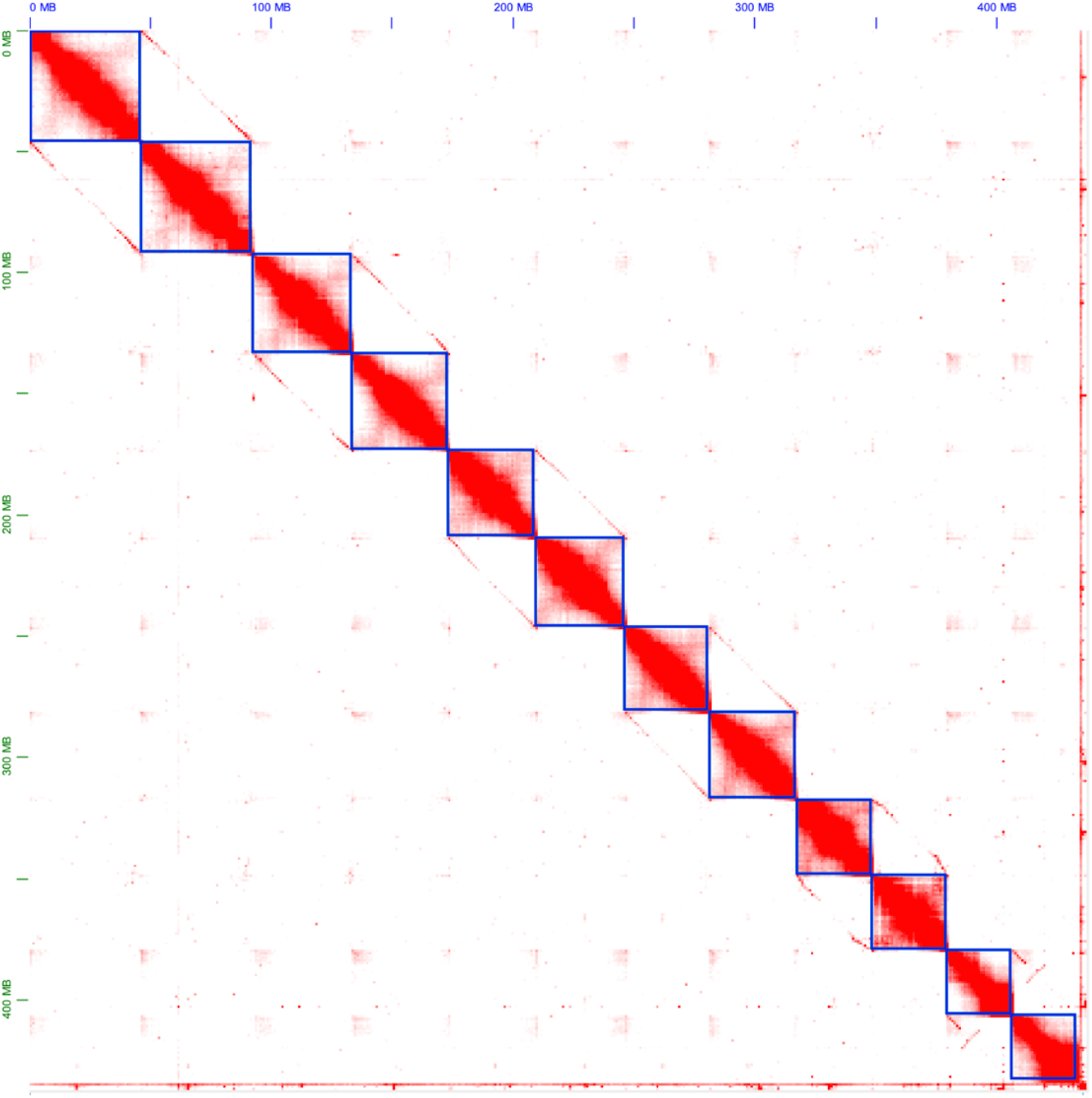
HiC contacts for the *Oxalis stricta* genome visualized by Juicebox (Durand *et al*. 2016a). Diagonal interactions indicate relationships between blue boxes (pseudomolecules) which show subgenome similarity.

### Genome annotation

*De novo* annotation of repetitive sequences followed by repeat masking revealed 39.7% of the *O. stricta* genome was repetitive with retroelements and unclassified interspersed repeats composing most of the repetitive sequence (Fig 3, Supplementary Table 6). Annotation of protein-coding genes revealed 61,550 representative high confidence genes that encoded 115,089 gene models attributable to the extensive RNA-seq and full-length cDNA sequences available for annotation; 121,961 working gene models were annotated (Supplementary Table 7). The subgenomes represent 99.7% of the gene models, with only a small fraction (0.3%) encoded on the unanchored scaffolds. BUSCO assessment of the representative high confidence gene model gene annotations closely matched the genome-level BUSCO assessment, with 96.3% complete BUSCOs that were mostly duplicated with very few missing BUSCO orthologs (Supplementary Table 5).

**Figure 3:**
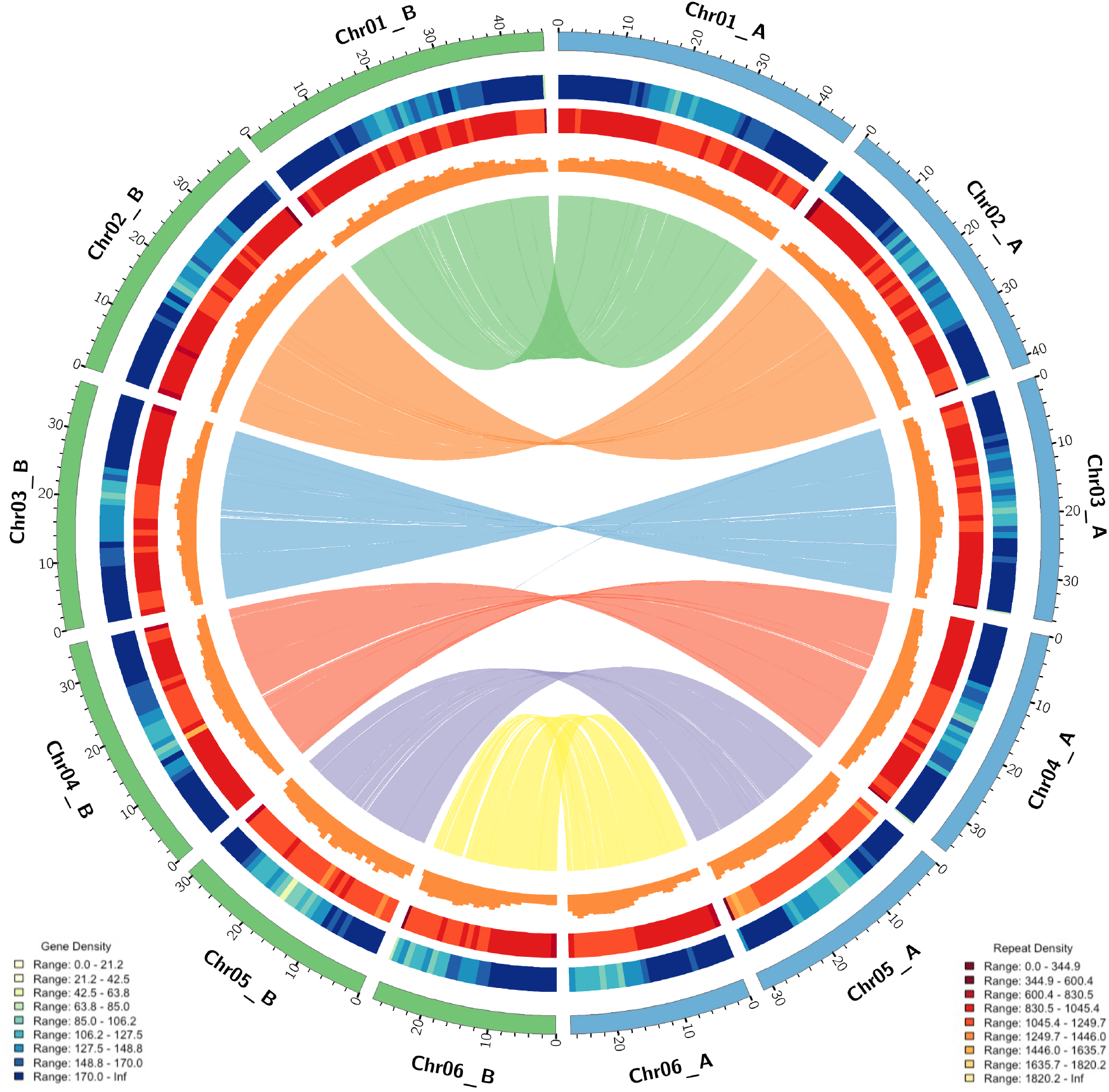
Circos plot (Krzywinski *et al*. 2009) of the *Oxalis stricta* genome. Rings (outermost to innermost) represent the 12 chromosomes binned into subgenomes A (blue) or B (green) in Mb, heatmap of gene density, total repeat density heatmap, histogram of transposable element density, and synteny between the two subgenomes. Heatmap and histogram bins are 1Mb in size.

### *O. stricta* is an allotetraploid

Previous reports indicate that *O. stricta* can occur as a tetraploid (Vaio *et al*. 2013). This is consistent with the pairing of homeologous chromosomes observed in our Hi-C signal, which reveals the presence of two subgenomes within our assembly, as demonstrated by diagonal similarities in the contact map (Fig. 2). Using sequence similarity of the genes to *A. carambola*, we assigned each of the 12 chromosomes to subgenome A or subgenome B (Fig. 3). Examination of each subgenome separately showed high percentage of single copy BUSCO’s with 95.5% and 96.1% for subgenome A and B, respectively (Supplementary Table 5). Assessing the subgenomes independently revealed a mere 3.7% and 4.3% duplicated BUSCO’s present in subgenome A and B, respectively. This reduction in duplicated BUSCOs in the full genome versus subgenome level analysis suggests that the subgenome assignment did not place two of the same chromosomes within one given subgenome. Distribution of genes across the two subgenomes was also even, with subgenome A containing 30,611 representative high confidence genes and subgenome B containing 30,667 representative high confidence genes.

### Repetitive sequences in *O. stricta*

To further annotate transposable elements within *O. stricta*, we used the EDTA software revealing 101.5 Mb of transposable element sequences accounting for 23.24% of the assembly (Supplementary Table 8, Fig 3). Of the different transposable element classes, there were 15.7% (68.5 Mb) and 9.6% (32 Mb) of Class I and Class II transposable elements, respectively. The Class I TEs were comprised of 3.2% (13.8 Mb) LINE elements and 12.5% (54.7 Mb) LTR elements. The Class II TEs were comprised of 5.9% (25.9 Mb) TIR elements and 1.4% (6 Mb) Helitron elements. Transposable elements can differ among subgenomes in an allopolyploid (Hosaka *et al*. 2024) and subgenome A contained 49.5 Mb (22.8%) of transposable elements while subgenome B contained 46.3Mb (21.42%) of transposable elements, showing a slight difference between the two subgenomes. While differences in individual transposable element classes were minimal between the two subgenomes, the three largest differences (>0.5% difference between subgenomes) were for *Copia* and *Mutator* elements that were more prevalent in subgenome A while unknown LTR elements were more prevalent in subgenome B. *Jockey* elements were present only within subgenome A with 129 elements representing 36 kb.

### Orthology analyses reveal lineage-specific genes within *O. stricta*

OrthoFinder2 was used to build a phylogeny for *O. stricta* that was consistent with the known phylogenetic relationships of these species (Fig. 4A). *C. follicularis*, a member of the Oxalidales, was correctly placed sister to the three Oxalidaceae species in the phylogeny. The two *Oxalis* species were also placed sister to one another. Clustering of 276,072 proteins from these seven species resulted in 27,532 orthogroups. Of these, 221 orthogroups were specific to the Oxalidales, 354 orthogroups were specific to the Oxalidaceae, 303 orthogroups were specific to the *Oxalis* genus and 2,764 orthogroups were unique to *O. stricta* which was comprised of 8,277 genes (Fig. 4B). The *O. stricta* specific genes were enriched for the biological process GO terms GO:0035194 (regulatory ncRNA-mediated post-transcriptional gene silencing; p-value 0.00021), GO:0019941 (modification-dependent protein catabolic process; p-value 0.00535), GO:0006266 (DNA ligation; p-value 0.01876), GO:0006273 (lagging strand elongation; p-value 0.01876), GO:0006310 (DNA recombination; p-value 0.02161) and GO:0050832 (defense response to fungus; p-value 0.02161) (Supplementary Fig. 2). Within *O. stricta* there were orthogroups specific to each subgenome, with subgenome A containing 504 exclusive orthogroups and subgenome B containing 452 exclusive orthogroups. The subgenome A-exclusive orthogroups contained 1,289 genes while the subgenome B-exclusive orthogroups contained 1,090 genes.

**Figure 4:**
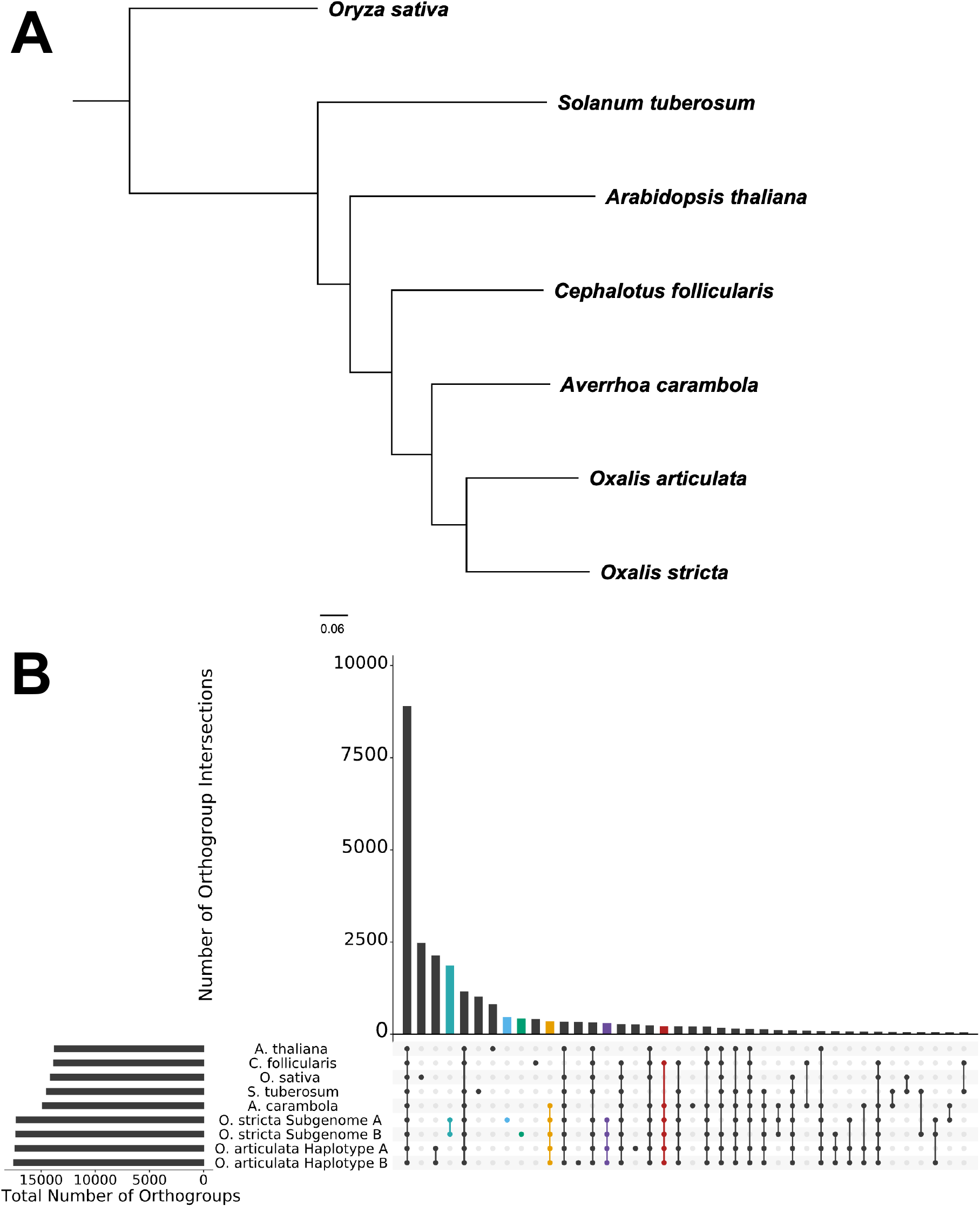
Phylogeny and orthology of *Oxalis stricta*. A. Phylogeny of *O. stricta* and other angiosperms using Orthofinder2 (Emms and Kelly 2019). B. Upset plot of orthogroups identified between *Arabidopsis thaliana, Averrhoa carambola, Cephalotus follicularis, Oryza sativa, Oxalis stricta* and *Solanum tuberosum* using Orthofinder2. Light blue represents *O. stricta* subgenome A specific orthogroups, green represents *O. stricta* specific subgenome B orthogroups, teal represents *O. stricta* subgenome shared orthogroups, purple represents *Oxalis* specific orthogroups, and orange represents Oxalidaceae specific orthogroups, and red represents Oxalidales specific orthogroups.

### *O. stricta* subgenomes are highly syntenic

Synteny analysis between *O. stricta* subgenomes A and B revealed 91 blocks containing 40,705 syntenic genes spanning across 211 Mb and 210 Mb in subgenome A and B, respectively (Supplementary Table 9, Figure 3). Examination of the syntenic relationships revealed that a majority of syntelogs exist within a 1:1 relationship between the two subgenomes. A total of 15,599 1:1 syntelogs were present, accounting for 76.64% of the syntenic relationships between the two subgenomes. A total of 9,507 other genes had syntenic relationships, with the next most prevalent relationship being 2:2 between the subgenomes. These 2:2 syntelogs comprised 7,220 genes in total and represent instances of paralogous genes. As these paralogs are present in both subgenomes their duplication is likely ancestral to the most common ancestor of the two subgenomes. The next two most common relationships within the syntelogs were 1:2 and 2:1, representing 1,149 and 1,074 genes, respectively, that likely represent recent gene duplications or gene losses that uniquely occurred within one of the subgenomes.

### Synteny within the Oxalidaceae

We performed synteny analysis between the *O. stricta* subgenomes, *A. carambola* and the haplotypes of *O. articulata* (Fig. 5). A total of 352 syntenic blocks comprised of 29,160 genes were found between subgenome A and *A. carambola*, with similar synteny between *A. carambola* and *O. stricta* subgenome B observed (Supplementary Table 10). These syntenic genes were dominated by 1:1:1 syntelogs, totaling 10,577 triplets, between *O. stricta* subgenome A, *O. stricta* subgenome B and *A. carambola*. When comparing the subgenomes of *O. stricta* with the haplotypes of *O. articulata*, we found that there were more syntenic blocks and collinear gene content with haplotype A of *O. articulata* for both subgenomes of *O. stricta*. Surprisingly, the percentage of collinear genes between *O. stricta* and *O. articulata* was less than that between *O. stricta* and *A. carambola*. The *Oxalis* genus is large, spanning over 500 species (Azkue 2000), of which, *O. stricta* and *O. articulata* belong to different sections within the genus (Oberlander *et al*. 2009). This diversity may explain why *O. stricta* is more similar to star fruit than *O. articulata*. Synteny across all of the available Oxalidaceae genomes revealed 9,021 genes present in a 1:1:1:1:1 relationship.

**Figure 5:**
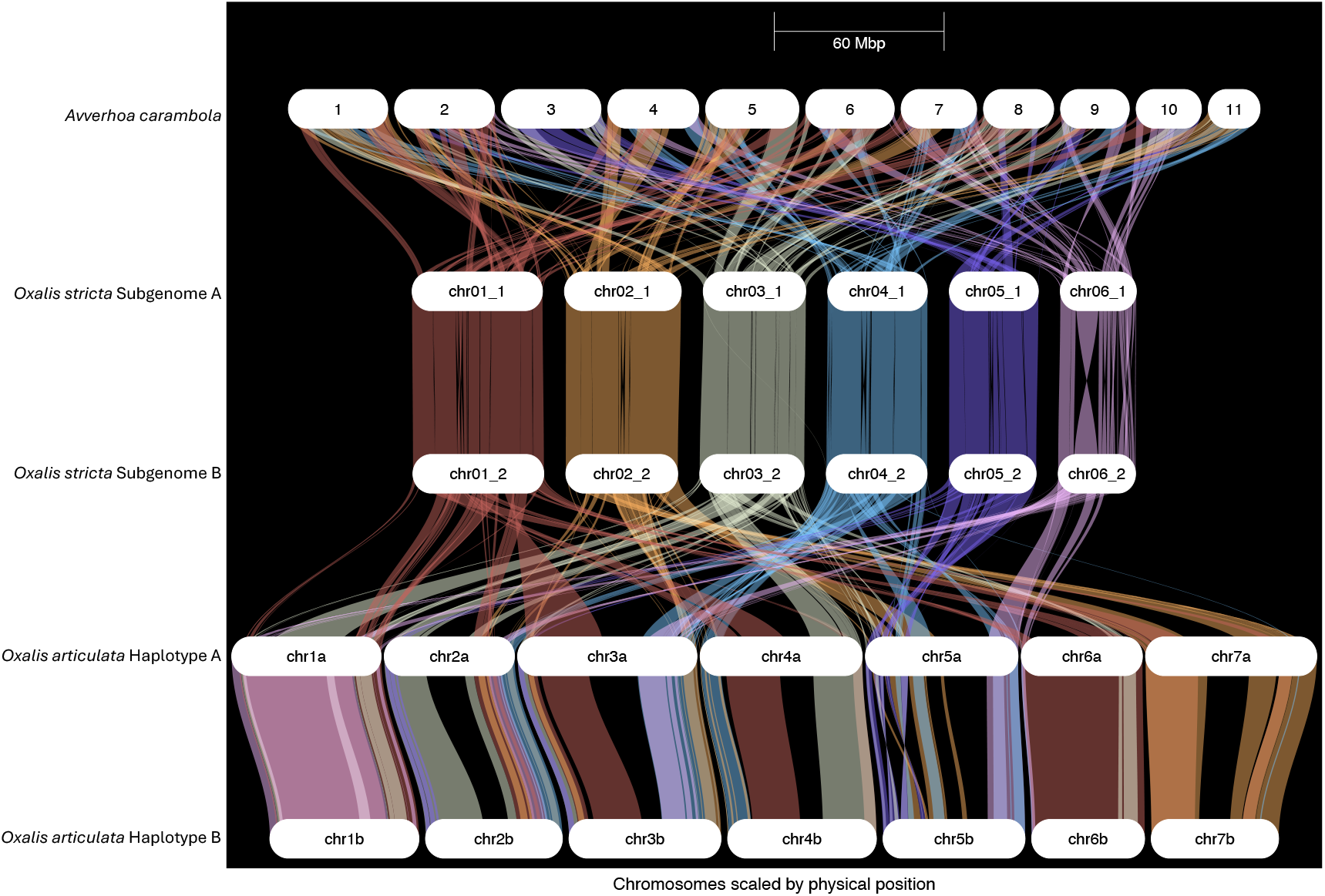
Synteny between the two *O. stricta* subgenomes, the two haplotypes of *O. articulata*, and *Averrhoa carambola.*

### Subgenome dominance in *O. stricta*

The 15,599 1:1 syntelogs between the two *O. stricta* subgenomes were examined for subgenome dominance using gene expression data from 13 replicated RNA-seq datasets comprising different tissues and/or treatments. Of these 1:1 syntelogs, a total of 30,040 of the genes had a transcript per million (TPM) above 0 in our expression profiling datasets. A Wilcoxon rank sum test of the mean log(TPM+1) values was conducted to test whether syntelogs within a given subgenome showed dominance in certain tissues or conditions, excluding the diurnal time-course due to it containing substantially more samples than the other tissue and treatment sets. Across the 12 datasets included, both flower (open and closed) and root (control and salt treated) tissues showed a bias, in which the median TPM’s are higher in subgenome B than subgenome A (Fig. 6A, Supplementary Figure 3). Syntelog pairs were defined as biased if the log2 expression fold difference between the subgenomes was greater than |2|. Examination of the paired syntelog expression within the flower tissues further showed overall bias for subgenome B, with a total of 2,214 pairs in closed flowers and 2,161 pairs in open flowers having bias for subgenome B (Supplementary Table 11). Examination of the paired syntelog expression within the root tissues also revealed overall bias for subgenome B, with a total of 2,125 pairs in control roots and 2,251 pairs in salt treated roots (Supplementary Table 12).

**Figure 6:**
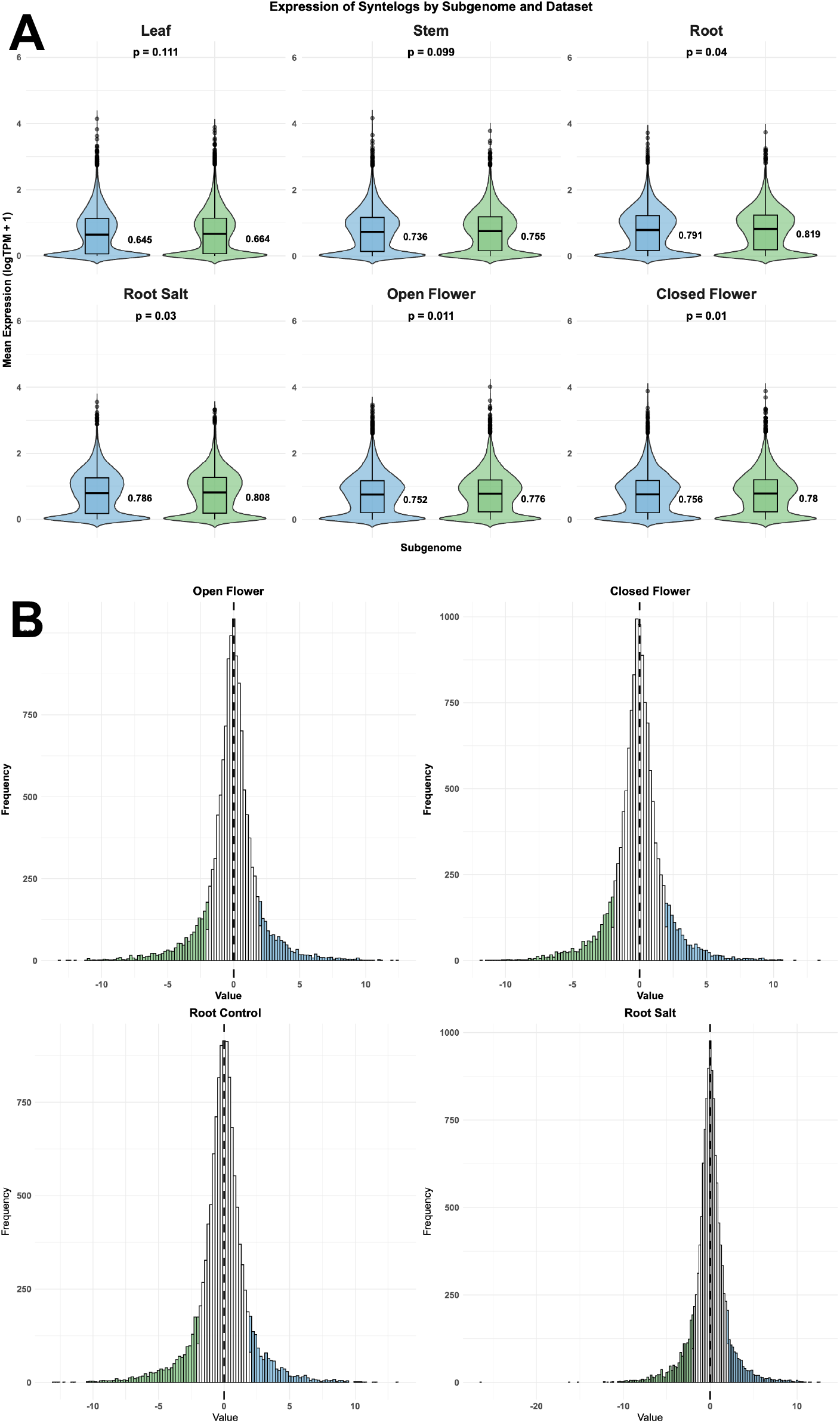
Expression of 1:1 syntelogs in *O. stricta* and their expression bias. A. Violin plots depicting expression of 1:1 *O. stricta* subgenome syntelogs in various tissues/treatments. P-values above each subgenome comparison are the pair-wise Wilcoxon test assessing for significant differences between the subgenomes. Values beside boxplots are median log(TPM+1) expression values for each subgenome/dataset. Subgenome A in blue and subgenome B in green. B. 1:1 syntelog expression within flower tissues and root tissues. Syntelogs showing bias, a log2 expression fold difference of |2|, are labeled in blue (subgenome A) or green (subgenome B) (Supplementary Table 9).

Within the set of closed flower subgenome B biased genes, genes annotated to be involved in lipids, seed storage and ripening were present. For example, the most biased gene for closed flowers was orthologous to *AtCLO1/AtCLO2/AtCLO3* which encode for caleosin proteins implicated in the pollen coat in some species (Hanano *et al*. 2023). Interestingly, the homolog of PHYTOENE DESATURASE 1 (*AtPDS1*) was also biased for the subgenome B syntelog in both open and closed flowers. *PDS1* is vital for the production of plastoquinone and tocopherols (Norris *et al*. 1995). Plastoquinone is a key component of the electron transport chain in chloroplasts and acts as a cofactor for PHYTOENE DESATURASE 3 (*PDS3*) which is involved in carotenoid biosynthesis. This availability of plastoquinone may be important in the floral tissues of *O. stricta* as the flower petals are yellow indicative of carotenoids. Both plastoquinone and tocopherols are also important protectors from reactive oxygen species and may help protect the floral organs from stress.

Within the set of root subgenome B biases genes, genes involved in transport, DNA binding and plant defense were present. Examples of the most biased genes in the control roots included Oxst.03_2G0008650, annotated as a ABC-2 type transporter and Oxst.01_2G027370, annotated as a PADRE domain containing protein, which are linked to plant defense against fungal pathogens and abiotic stress (Didelon *et al*. 2020). Within the salt treated roots, we observed that Oxst.03_2G052820 had a high expression bias of -15.22 for the subgenome B syntelog, which was annotated as a senescence/dehydration-associated protein. These analyses highlight that subgenome bias is limited within *O. stricta* and restricted to flower and root tissues. However, these biased genes may play important roles in reproduction and responses to the environment within these organs.

Subgenome dominance, a phenomenon in which one parental subgenome in a polyploid exhibits greater gene expression abundance than other subgenome(s), is hypothesized to be influenced by the degree of similarity between the constituent subgenomes (Garsmeur *et al*. 2014). Previous studies suggest that subgenome dominance is less pronounced or prevalent in autopolyploids (derived from the duplication of a single genome) or in allopolyploids formed from species with highly similar subgenomes (Garsmeur *et al*. 2014; Zhao *et al*. 2017; Alger and Edger 2020; Fang *et al*. 2024). This is thought to be because the greater the similarity between subgenomes, the fewer genetic conflicts may arise from their co-existence in a single nucleus, thus reducing the need for differential regulation or gene silencing. In our *O. stricta* assembly, the two subgenomes exhibit a high degree of similarity, with over 60% of their genes being collinear, indicating a close evolutionary relationship. Consequently, the general lack of significant subgenome dominance observed across 8 of our 12 datasets aligns with this hypothesis, suggesting that the close relatedness of the *O. stricta* subgenomes has limited the establishment of strong expression biases.

## CONCLUSIONS

Here, we present a chromosome-scale genome assembly for allotetraploid yellow wood sorrel (*O. stricta*). A comparative analysis of the two subgenomes revealed the presence of large syntenic blocks and a high degree of collinearity, with over 60% of the genes occupying corresponding positions. Furthermore, synteny analysis comparing the yellow wood sorrel subgenomes with the only two other available genomes within the Oxalidaceae, star fruit (*Averrhoa carambola*) and *O. articulata*, also showed high percentages (50% and 47%, respectively) of collinear genes, highlighting conserved genomic regions within the family. Subgenome dominance analysis between syntenic gene pairs revealed some tissue-specific expression bias, with flower and root tissues exhibiting favored expression for subgenome B, while the remaining tissues and treatments showed no significant bias in expression between the two subgenomes. This newly generated genome assembly represents only the third publicly available genome in the Oxalidaceae, and second within the large *Oxalis* genus, and will provide a valuable resource for future research into the evolutionary and functional genomics of this diverse plant group.

## Abbreviations

BLAST: Basic Local Alignment Search Tool
bp: base pairs
benzothiadiazole: BTH
BUSCO: Benchmarking Universal Single-Copy Orthologs
cDNA: complementary DNA
CRL: custom repeat library
Gb: gigabase pairs
kb: kilobase pairs
LAI: LTR Assembly Index
LTR: long terminal repeat
Mb: megabase pairs
Methyl Jasmonate: MeJA
MF: Molecular function
mRNA: messenger RNA
NCBI: National Center for Biotechnology Information
nt: nucleotide
ONT: Oxford Nanopore Technologies
PASA: Program to Assemble Spliced Alignments
RNA-Seq: RNA-sequencing
ROS: Reactive Oxygen Species
SRA: Sequence Read Archive
TPM: transcripts per million

## DATA AVAILABILITY

Raw sequence data have been deposited in the National Center for Biotechnology Sequence Read Archive under BioProject PRJNA856298.

## ACKNOWLEDGEMENTS

We thank the Michigan State University Research Technology Support Facility and the Texas A&M AgriLife Research: Genomics and Bioinformatics Service for providing sequencing services.

## SUPPLEMENTARY FILES

Supplementary Table 1: Sequence datasets used in this study.

Supplementary Table 2: Oxford Nanopore Technologies whole genome shotgun sequence reads used in the *Oxalis stricta* assembly.

Supplementary Table 3: Assembly statistics for the *Oxalis stricta* assembly.

Supplementary Table 4: Pseudomolecule lengths and gap content for the *Oxalis strica* Assembly.

Supplementary Table 5: Benchmarking universal single copy orthologs (BUSCO) results on the *Oxalis strica* assembly and annotation.

Supplementary Table 6: Repetitive sequence content in the *Oxalis stricta* genome assembly.

Supplementary Table 7: *Oxalis stricta* gene annotation summary.

Supplementary Table 8: Transposable elements in *Oxalis stricta*.

Supplementary Table 9: *Oxalis stricta* synteny results.

Supplementary Table 10: *Oxalis stricta, O. articulata* and *Averrhoa carambola* GENESPACE pangene results.

Supplementary Table 11: *Oxalis stricta* flower subgenome B dominant genes.

Supplementary Table 12: *Oxalis stricta* root subgenome B dominant genes.

Supplementary Figure 1: K-mer plot of the *Oxalis stricta* genome produced by the KAT program.

Supplementary Figure 2: GO (Gene Ontology) terms enriched for *O. stricta* specific genes identified by OrthoFinder.

Supplementary Figure 3: Expression of 1:1 syntelogs in *O. stricta* leaf tissue treatments and their expression bias.

## AUTHOR’S CONTRIBUTIONS

CRB and PPE designed the study. PPE obtained germplasm. JB, JCW generated data. BV, JPH, and JCW performed data analyses. JCW and CRB wrote the manuscript; all coauthors reviewed and edited the manuscript.

