## Supplementary Figures for "Chromosome-scale genome assembly for Yellow Wood sorrel, *Oxalis stricta*"

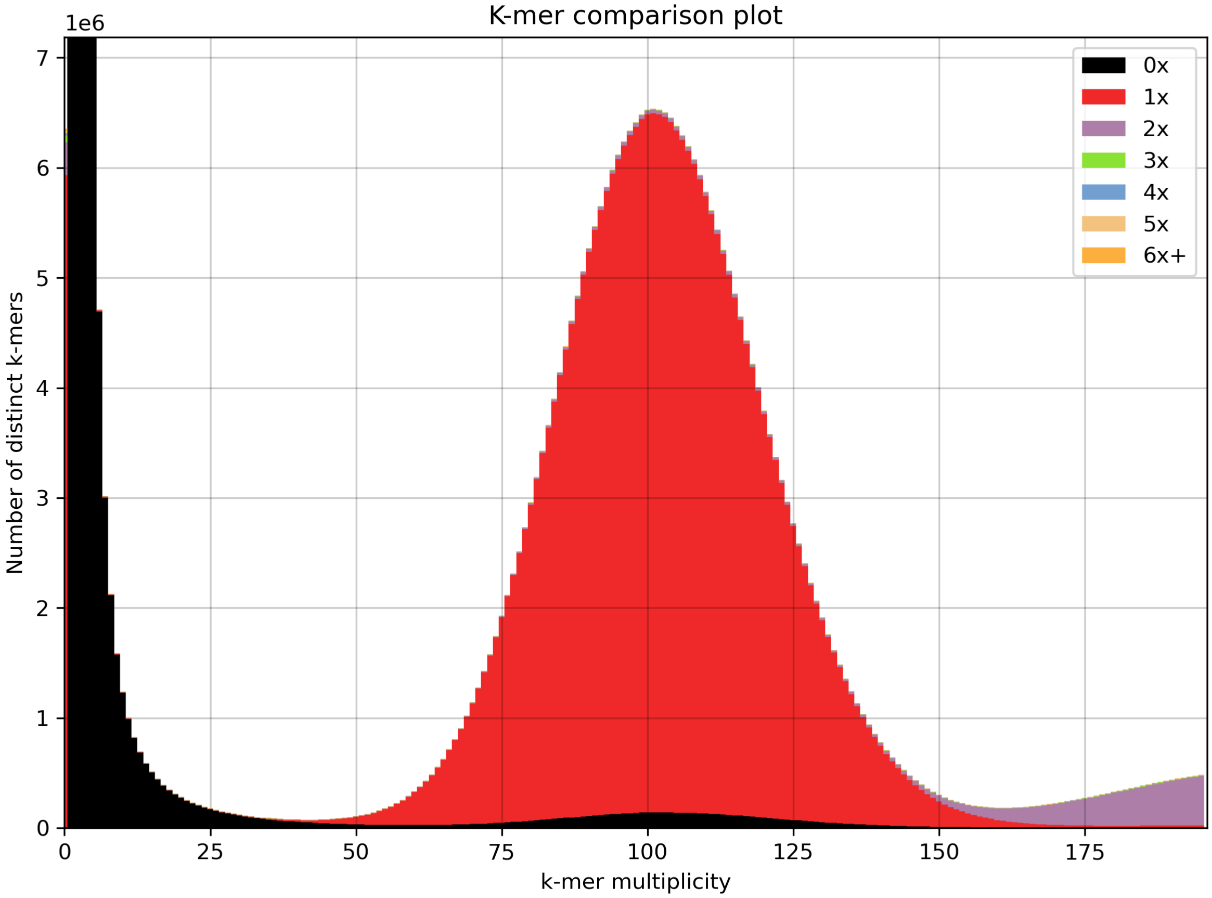


Supplementary Fig. 1: K-mer plot (k = 21) of the *Oxalis stricta* genome produced by the KAT program (Mapleson et al. 2017).

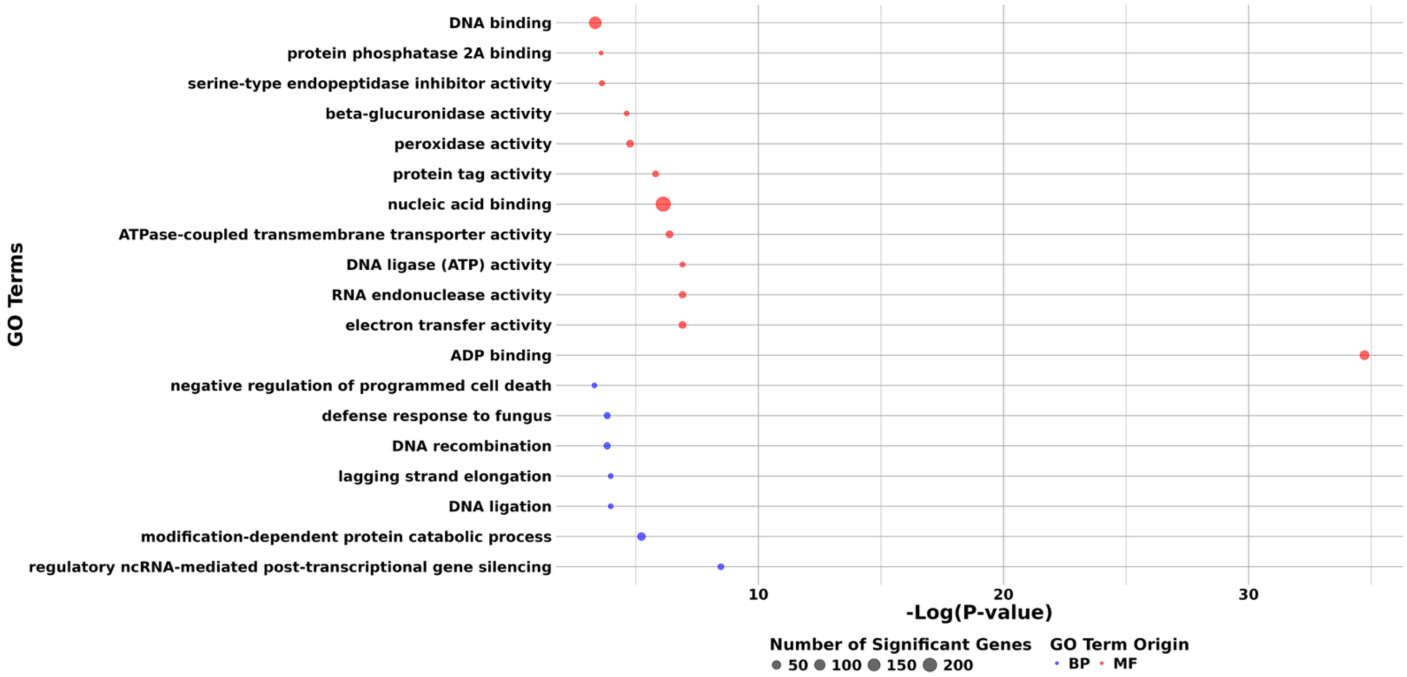


Supplementary Fig. 2: GO (Gene Ontology) terms enriched for *O. stricta* specific genes identified by OrthoFinder. BP = Biological Process, MF = Molecular Function

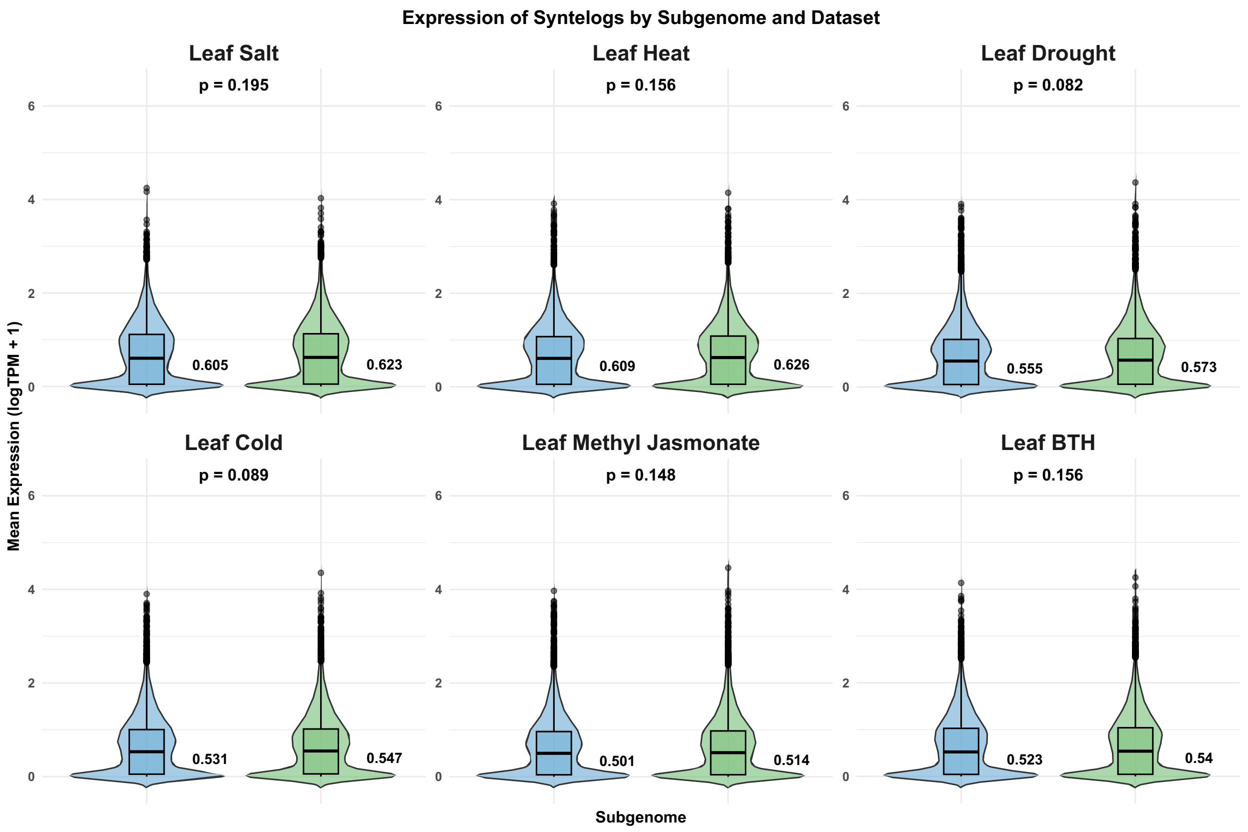


Supplementary Fig. 3: Expression of 1:1 syntelogs in *O. stricta* leaf tissue treatments and their expression bias. Violin plots depicting expression of 1:1 *O. stricta* subgenome syntelogs in leaf treatments. P-values above each subgenome comparison are the pair-wise Wilcoxon test assessing for significant differences between the subgenomes. Values beside boxplots are median log(TPM+1) expression values for each subgenome/dataset. Subgenome A in blue and Subgenome B in green.
